# Hypoxia-mediated fine-tuning of the TLR7/9-triggered human PDC-derived IFN-α response is mediated by combined cellular and soluble IFN-regulators

**DOI:** 10.1101/2024.09.15.613102

**Authors:** Alexander Lenkewitz, Ibtissem Ben Brahim, Felix Herrmann, Noor-A-Kasida Islam, Olga Ticha, Isabelle Bekeredjian-Ding

**Author notes:** Correspondence: Prof. Dr. med. Isabelle Bekeredjian-Ding, Institute of Medical Microbiology and Hospital Hygiene Philipps-University of Marburg, Hans-Meerweinstr. 2, D-35043 Marburg, Germany, Mail Fon. This study includes data from the PhD thesis of A.L and I.B.B.

## Abstract

Hypoxia occurs in settings where stringent control of inflammation is mandatory to avoid tissue damage. Here, we analyzed how hypoxia-induced mediators alter recognition of pathogen-derived molecular patterns (PAMPs). The most relevant finding was that hypoxia selectively targets IFN-α induced upon stimulation with nucleic acid-sensing Toll-like receptors (TLR7/8 and -9) in PBMC. Notably, IFN-α secretion was reduced, but IFN-α producing capacity preserved. Corroborating these findings IFN-dependent cytokines IP-10 and IL-12 p70 were reduced under hypoxia, while other cytokines such as TNF remained unaffected. A role for hypoxia-inducible factors (HIF) was confirmed with prolyl hydroxylase (PHD) inhibitors CoCl_2_ and DMOG. Plasmacytoid dendritic cells (PDC) were identified as the major source of IFN-α and experiments in B cell/PDC and monocyte/PDC cocultures indicated that T cells are not required and both lymphoid and myeloid cells equally support inhibition of IFN-α. Moreover, supernatants from unstimulated PBMC generated under hypoxic conditions were not suppressive but further experiments indicated that hypoxia-specific release of IFN-α regulators lactate and TGF-β can contribute to the IFN-α blockage and might synergize with PGE_2_. However, each one of these factors alone is insufficient to block IFN-α secretion. These findings suggest that control of the PDC response occurs via combined soluble factors and cell contact-dependent mechanisms, thereby enabling a fine-tuning of the anti-microbial immune response in hypoxia.

## Introduction

Hypoxia is a permanent condition in specific tissues, such as cartilages or light zones of the germinal centers [1, 2] but can also arise in inflamed or infected tissues [3]. Cellular sensing of oxygen is based on oxygen-dependent stability of hetero-dimeric transcription factors termed hypoxia-inducible factors (HIF) that regulate the transcriptional activity of hundreds of genes [4–8]. The transcriptional changes affect fundamental pathways, most prominently a metabolic switch from oxidative respiration to glycolysis, resulting in the secretion of lactate and H^+^ [9].

Notably, cell culture conditions mimic essential aspects of *in vivo* conditions including temperature and CO_2_ concentration. However, oxygen concentrations under standard cell culture conditions reach approximately 18.6% [10, 11] albeit oxygen concentrations *in vivo* range between 1-11% [12]. Potential inaccuracies that arise from this deviation were discussed in [11] and effects recently reviewed in [13]. Specific aspects of the impact of hypoxia on innate immune cell functions were addressed in [14–18].

Both hypoxia and the recognition of bacteria via PAMPs (TLR4 [19], TLR2 [20]) induce signaling via Hypoxia-inducible factors (HIF) and NFкB, a key transcription factor enabling pro-inflammatory immune responses. Thus, hypoxia and inflammatory cell signaling act in concert, resulting in enhanced innate immune cell functions and bacterial clearance [15, 21, 22]. Investigation of the impact of hypoxia upon T cell mitogen exposure demonstrated increased levels of IL-2, IL-4, IFNγ, IL-6 and reduced amounts of IL-10 after stimulation of PBMC [23, 24] and conflicting results after T cell stimulation with anti-CD3/CD28 [25–28]. Furthermore, hypoxia increased expression of TNF in murine macrophages and human monocytes [29, 30]. Previously, lactate, an hypoxia-specific metabolite was shown to reduce secretion of pro-inflammatory cytokines. This was demonstrated upon stimulation with LPS in murine macrophages (IL-12 p40, IL-1β and IL-6) [31, 32], human PBMC treated with 15 mM lactate (IL-1β upon simultaneous stimulation with ATP) [31] and with 3.5 or 15 mM lactate (TNF, IL-1β, IL-6 and IL-10) in [33], in human monocytes (TNF expression) [34], and monocyte-derived dendritic cells exposed to lactic acid (IL-12, TNF, IL-6) [35].

Altogether, hypoxia-mediated modulation of the immune response is complex. The currently available data provide an incomplete understanding and contradictory results. It, therefore, remains highly relevant to investigate the effects of hypoxia on immune cell function. Here, we sought to identify factors regulating the reactivity of human leukocyte response under hypoxic conditions. Most prominently, we identified control of IFN-α production as a major factor in hypoxia-mediated modulation of the human immune response.

## Materials and methods

### Ethics statement

The use of PBMC isolated from buffy coats was approved by the ethics committee of the Medical Faculty of the University of Frankfurt (approval #154/15).

### Cell culture

Cells were incubated in normoxia (air + 5% CO_2_, 80% humidity, 37°C) or hypoxia (1% O_2_, 5% CO_2_, 80% humidity, 37°C) monitored by the oxygen sensor of the InVivO_2_ 400 hypoxic chamber (Baker, Bridgend, UK) or the M55 HEPA variable atmosphere workstation (Don Whitley Scientific Limited, Bingley, UK) used for co-culture experiments in Fig. 4 and one PDC experiment in Fig. 3c). For cell culture medium RPMI (Thermo Fisher Scientific, Dreieich, Germany) was supplemented with 10% heat inactivated (56°C for 60 min) fetal calf serum (Sigma Aldrich, Munich, Germany), 1% 100 U/ml penicillin + 10mg/ml Streptomycin solution (Biochrom AG, Berlin, Germany), 1% 2 mM L-glutamine (Biochrom AG, Berlin, Germany) and 1% 1M HEPES pH 7.4 (own production). Media were adapted to hypoxia for 4 hours.

### Cell isolation

PBMC were isolated from buffy coats (German Red Cross South Transfusion Center in Frankfurt am Main, Germany) via gradient centrifugation with Pancoll (Pan BioTech, Aidenbach, Germany). Erythrocytes were lysed with 0.86% ammonium chloride. Cell numbers and viability were measured with trypan blue (Gibco by Life Science, Darmstadt, Germany) in a hemocytometer (TC20 Automated Cell Counter by Bio-Rad Laboratories, Feldkirchen, Germany). PBMC were adapted to culture conditions (normoxia, hypoxia, lactic acid) for 4 hours at 4x10^5^ cells / well in U-well 96-well plates and subsequently stimulated for 20 hours. Supernatants were frozen at -20°C and cells stained for flow cytometry as indicated. PBMC supernatants for PDC stimulation were incubated as described but left unstimulated. After harvest they were frozen at -150°C until further use.

PDC were depleted from PBMC by AutoMACS (deplete_s) after preincubation with Fc block and anti-CD304 microbeads (BDCA-4/Neuropilin-1) MicroBead kit (Miltenyi Biotec, Bergisch-Gladbach, Germany). Total PBMC controls were subject to the same AUTOMACS protocol. PDC purity was controlled by flow cytometric analysis: the mean PDC frequency after depletion was 0.10% (SEM ±0.013) versus 0.33% (SEM ±0.051) in total PBMC (Fig. S3a). PBMC and PDC-depleted PBMC were adapted to culture conditions for 4 hours before stimulation at 4x10^5^ cells / well in U-well 96-well plates for 20 hours. Supernatants were frozen at -20°C. PDC from different donors were isolated as described and exposed to thawed PBMC supernatants for 4 hours in V-shaped wells of 96-well plates before stimulation. Supernatants were frozen at -20°C.

Leukocyte subpopulations were isolated from PBMC. PDC were isolated via autoMACS or MidiMACS using LS Columns (Miltenyi Biotec, Bergisch-Gladbach, Germany), with the anti-CD304 (BDCA-4/Neuropilin-1) MicroBead kit (Miltenyi Biotec, Bergisch-Gladbach, Germany) and identified as BDCA-2 positive, CD19 negative cells via flow cytometry. PDC purity ranged between 85.3 and 98.25%. Enriched PDC were adapted to culture conditions (normoxia, hypoxia, 10 mM K^+^ lactate) for 4 hours before stimulation in V-well 96-well plates for 20 hours (Fig. 3c) or 18 ±4 hours (Fig. 5c) at 5x10^4^ cells/well.

For co-cultures B cells were isolated together with PDC by AutoMACS (possel_s program followed by possel) after combined incubation with Fc block, anti-CD19 and anti-BDCA-4 microbeads (Miltenyi Biotec, Bergisch-Gladbach, Germany). Monocytes were enriched in conjunction with PDC with Fc block, anti-CD14 and anti-BDCA-4 microbeads and subjected to AutoMACS selection (possel_s program followed by possel) (Miltenyi Biotec, Bergisch-Gladbach, Germany). Purity of isolated cells and ratios (PDC/B cells; PDC/Monocytes) were controlled by flow cytometry: the mean frequencies were 2.24% PDC (SEM ±0.55)/ 93.80% B cells (SEM ±1.10) and 2.03% PDC (SEM ±1.20)/ 98,44% Monocytes (SEM ±0.36).

### Cell stimulation

Cells were treated with the following reagents purchased from Sigma-Aldrich, Munich, Germany: 1 µM SB431542 TGF-β RI Kinase Inhibitor VI (dissolved in DMSO), 10 mM L-Lactic acid (dissolved in dH_2_O), 10 mM K^+^ L-lactate (dissolved in dH_2_O), 20 µM Forskolin (dissolved in DMSO), *Escherichia coli* LPS (O111:B4). Following reagents were purchased from Merck, Darmstadt, Germany: 100 µM dimethyloxalylglycine (DMOG, dissolved in dH_2_O), 200 µM CoCl_2_ (dissolved in dH_2_O). DMOG and CoCl_2_ were titrated to a concentration in which they did not reduce the viability of PBMC within 20 hours (Fig. S2). Further stimulatory reagents used: 0.25 or 1 µg/ml Resiquimod (R848; InVivoGen, Toulouse, France, dissolved in PBS), 0.5 mM loxoribine (dissolved in dH_2_O; Sigma, Munich, Germany), 0.5 µM CpG2216 (5’-ggGGGACGATCGTCgggggG-3’, small letter nucleotides represent phosphorothioate modifications; Eurofins, Ebersberg, Germany, dissolved in dH_2_O).

### Flow cytometry

Cell viability was determined using LIVE/DEAD Fixable Aqua Dead Cell Stain Kit (Thermo Fisher Scientific, Dreieich, Germany) for 30 min. Cells were stained with fluorescence-labeled antibodies for 30 min at 4°C in PBS + 0.5% heat inactivated FCS. Measurements were performed on an LSRII SORP or an LSR Fortessa flow cytometer (BD Biosciences, Heidelberg, Germany). Data were analyzed with Kaluza

#### 2.1 (Beckman Coulter, Krefeld, Germany)

The following antibodies were used for flow cytometry: Anti-CD303 (BDCA-2)-FITC, clone 201A (Biolegend, San Diego, USA), anti-CD14-V450, clone MDP9, anti-CD19- APC, clone HIB19, (all from BD Biosciences, Heidelberg, Germany), anti-CD14- PE/Cy5, clone M5E2, (Biolegend, San Diego, USA), anti-CD19-PE/Cy7, clone J3- 119, (Beckman Coulter, Krefeld, Germany), anti-CD3-AF700, clone OKT3, (Biolegend, San Diego, USA).

### Cytokine measurements

Unless stated otherwise, cells were stimulated for 20 hours and cell-free supernatants harvested and frozen for further use. The quantification of cytokine release in the supernatants was determined by ELISA. The following kits were used according to the manufactureŕs recommendations: IFN-α matched antibody pairs kit (Thermo Fisher Scientific, Dreieich, Germany), BD OptEIA ELISA kit IP-10, IL-6, TNF, IL-12 p70, IL-1β (BD Biosciences, Heidelberg, Germany), PGE_2_ Multi-Format ELISA Kit (Arbor Assays, Ann Arbor, USA). Lactate concentrations were quantified using the L-lactate assay kit (Megazyme, Bray, Ireland). Absorbance was measured on a Tecan Sunrise absorbance reader (Tecan, Maennedorf, Switzerland). IL-10 was measured by MAGPIX multiplex assay on a Bio-Plex MAGPIX Multiplex reader (Bio-Rad Laboratories, Feldkirchen, Germany).

### HIF-1**α** Western Blot

PBMC were incubated in normoxia or hypoxia for 4 or 24 hours. After incubation, cells were washed twice with degassed, cold PBS. Cell pellets were frozen with dry ice and stored at -150°C. For lysis, cells were incubated on ice for 15 min with SDS lysis buffer (55.5 mM Tris-HCl [pH 6.8], 9% glycerol, 2.2% SDS). Before the lysis, the buffer was supplemented with Halt Protease inhibitor cocktail (Thermo Fisher Scientific, Dreieich, Germany). Samples were then sonicated twice for 10 seconds, 3 cycles each at 40% power by a Bandelin Sonopuls GM 2200 (BANDELIN, Berlin, Germany). Samples were clarified by centrifugation at 14,000 g for 10 minutes at 4°C. Supernatants were collected and frozen at -80°C. Protein concentrations were determined by BCA assay. 50 µg of each sample was loaded and run on 8% polyacrylamide gel at 90 V for 10 min, followed by 120 V for 90 minutes in a vertical electrophoresis cell (Bio-Rad Laboratories, Feldkirchen, Germany). Proteins were transferred to nitrocellulose membranes at 90 V for 90-100 minutes by a Mini Trans-Blot cell (Bio-Rad Laboratories, Feldkirchen, Germany). Membranes were blocked with TBST-5% dry milk at RT for 1 hour then incubated overnight with primary antibodies, rabbit anti-HIF-1α (HIF-1α polyclonal antibody 20960-1-AP, Protein tech, Planegg-Martinsried Germany) and murine anti-HDAC1 IgG ((10E2) mouse mAb, Cell Signaling, Leiden, The Netherlands) at 4°C. After washing, membranes were incubated with IRDye 800CW donkey anti-rabbit IgG and IRDye 680RD donkey anti-mouse IgG secondary antibody for 1 hour at RT in the dark and thereafter scanned on the IR imager Odyssey CLx and analyzed via Image Studio Lite 5.2.2 (LI-COR Biosciences, Bad Homburg, Germany).

### Statistical analyses

Statistical analysis was carried out with Prism 8.01 (Graphpad, La Jolla, USA). Data were either pairwise compared with two-tailed student’s t-test or Wilcoxon matched-pairs signed-rank test. P values are presented as: ns (not significant) p ≥ 0.05 * p < 0.05; ** p < 0.01; *** p < 0.005; **** p < 0.001.

## Results

### IFN-α and IP-10 levels are reduced after TLR stimulation of human PBMC under hypoxic conditions

To assess the effect of hypoxia on the innate immune response PBMC were stimulated with the TLR7/8 ligand R848 or the TLR4 ligand LPS, or a combination of both. PBMC viability was comparable under hypoxic and normoxic conditions (Fig. S1). Secretion levels of IL-1β, TNF, IL-6 and IL-12 p70 showed little to no alteration under hypoxic conditions, with a few exceptions: release of IL-1β and IL-6 was increased under hypoxic conditions when cells were stimulated with LPS alone (Fig. 1a). Secretion of IFN-α was reduced upon stimulation with 0.25 or 1 µg/ml R848 in combination with 10 ng/ml LPS or 1µg/ml R848 alone (Fig. 1b) and significantly reduced upon stimulation with 0.25 µg/ml R848 (Fig. 1c).Similarly, levels of IP-10 were reduced under the same conditions (Fig. 1d). Of note, combined stimulation with LPS and R848 synergistically increased IL-12 p70 secretion as previously reported [36] but was not significantly altered in hypoxia despite a clear trend for reduction of IL-12 p70 in the combined LPS/R848 condition.

**Fig. 1:**
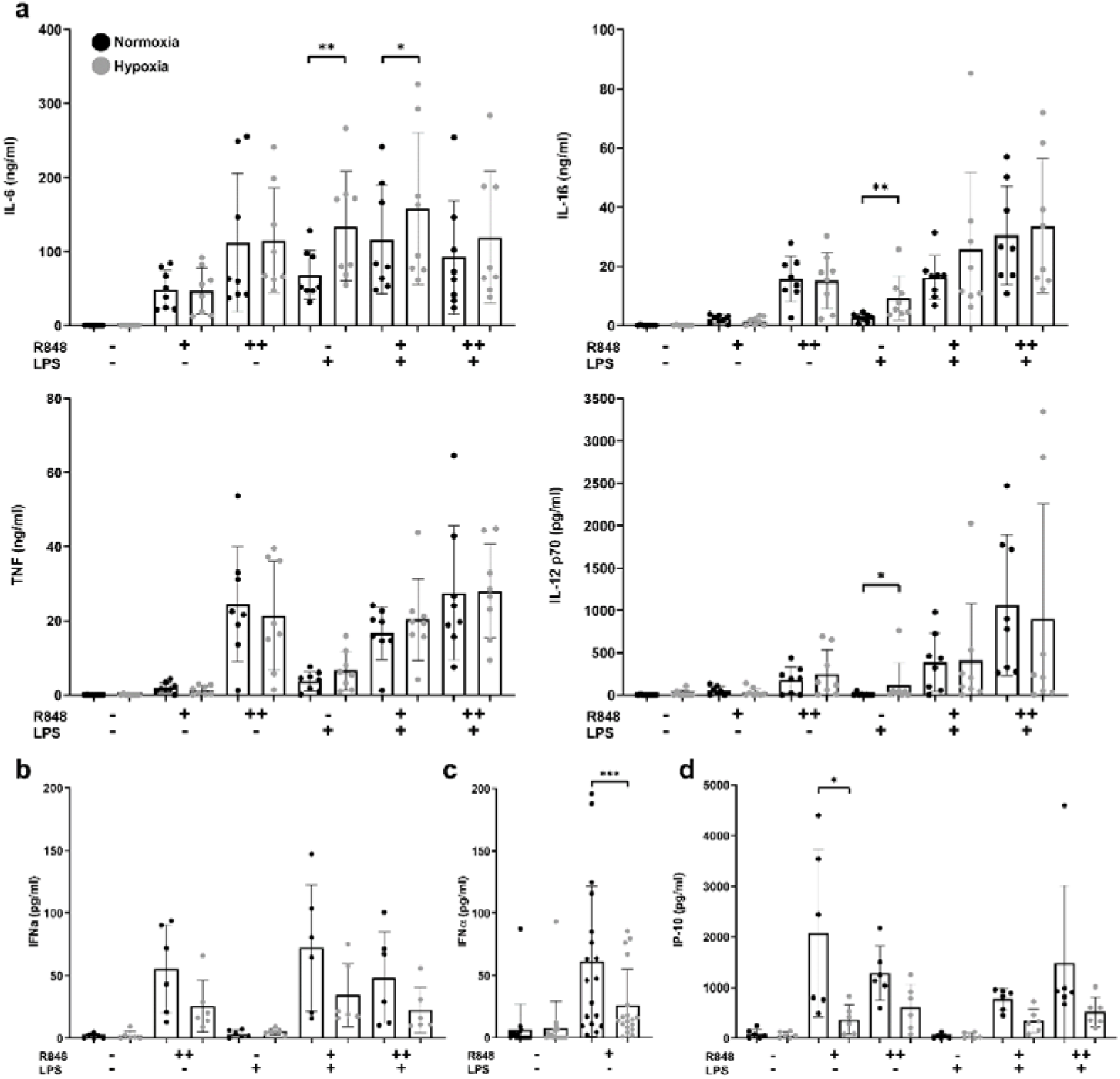
Cytokine secretion of PBMC under hypoxic conditions. PBMC were adapted to hypoxic conditions and stimulated with combinations of R848 and LPS: 0.25 (+) or 1 (++) µg/ml R848, 10 ng/ml *E. coli* LPS or both for 20 hours. Cytokines were measured via ELISA in supernatants. **a.** n = 8 donors. **b.** n = 6 donors; **c.** n = 18 donors; **d.** n = 6 donors; Bars indicate mean ± SD. Statistics: Wilcoxon matched-pairs signed-rank tests.

### PHD inhibitors mimic suppressive effects of hypoxia on release of IFN-**α**

Hypoxic culture conditions cause PHD inhibition, HIF stabilization and signaling. Controls of PBMC lysates via Western blot confirmed the presence of HIF-1α under hypoxic conditions and absence in normoxia (Fig. 2a). Hypoxia-mediated HIF stabilization can be mimicked *in vitro* by chemical inhibitors of PHD such as CoCl_2_ or DMOG. Here we used PHD inhibitors to assess a possible contribution of HIFs to suppression of IFN-α. To this end, PBMC preincubated with chemical inducers of hypoxia-related events were stimulated with R848, CpG2216 and loxoribine and IFN- α concentrations quantified in the supernatants (Fig. 2b). None of the PHD inhibitors caused a reduction in PBMC viability (Fig. S2). CoCl_2_ and DMOG severely reduced IFN-α secretion in response to loxoribine, a TLR7 agonist, and CpG2216, thus confirming a role for HIF. However, an IFN-α reduction after stimulation with R848 was only seen with CoCl_2_ and not observed for DMOG, despite only low IFN-α induction. Notably, lactate levels remained below the detection limit in all PHD inhibitor containing conditions (Fig 2c), thus ruling out a role for lactate in this setting.

**Fig. 2:**
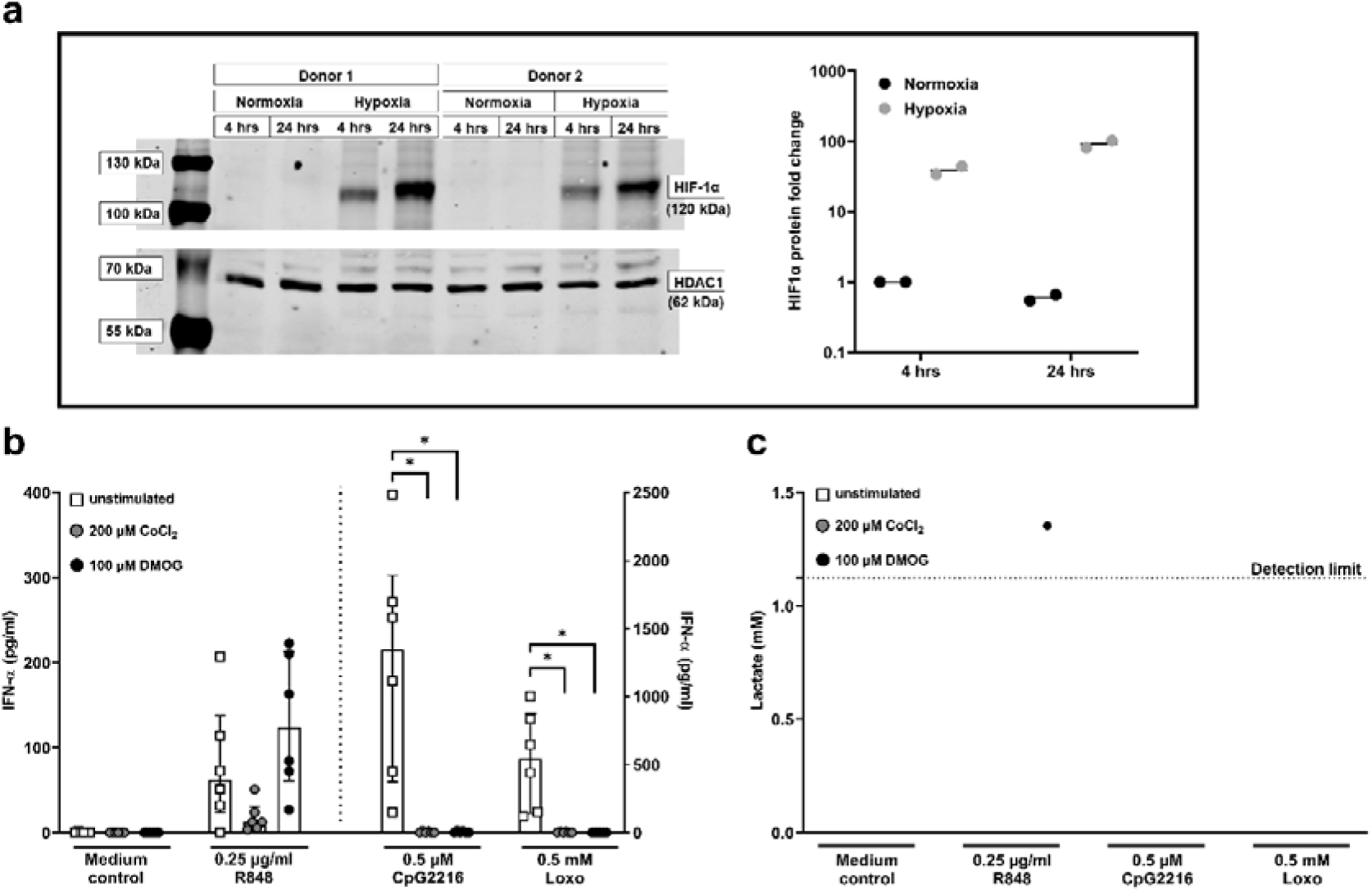
Hypoxia and PHD-inhibitor induced HIF-signaling and IFN-α reduction. **a**. PBMC were incubated in normoxia or hypoxia for 4 or 24 hours. Western blot controls of HIF-1α in PBMC whole cell lysates were performed as described. Western blots show band intensities of HDAC1 control and HIF-1α. The graph shows the fold- change of the HIF-1α signal in n = 2 donors. **b.** PBMC were exposed to PHD- inhibitors for 4 hours under normoxic conditions and then stimulated with 0.25 µg/ml R848 (left), 0.5 µM CpG2216 or 0.5 mM loxoribine (right) for 20 hours. IFN-α was measured in the supernatants. Three independent experiments were performed with n = 6 donors. Bars indicate median ± interquartile range calculated with Wilcoxon matched-pairs signed-rank test. **c**. Lactate was measured in the supernatants of PBMC exposed to PHD-inhibitors. Three independent experiments with n = 6 donors. The detection limit was 1.123 mM.

### PDC are the main IFN-**α** source and the target cell for hypoxia-mediated blockade of IFN-**α**

PDC are the most relevant professional human IFN-α secreting cells. They are characterized by high TLR7 and TLR9 expression [37] Since our results suggested that the hypoxia-induced IFN-α blockade might be promoted via selective inhibition of PDC-derived IFN-α production, we performed experiments with TLR ligands known to target PDC such as type A CpG oligonucleotide 2216 [38] and TLR7 agonist loxoribine [36, 39]. The results showed that induction of IFN-α by these ligands was blocked by exposure to hypoxic conditions. Again, this was accompanied by reduction of IP-10, an IFN-dependent cytokine (Fig. 3a), confirming that loss of IFN-α affects downstream effects of IFN-I.

**Fig. 3:**
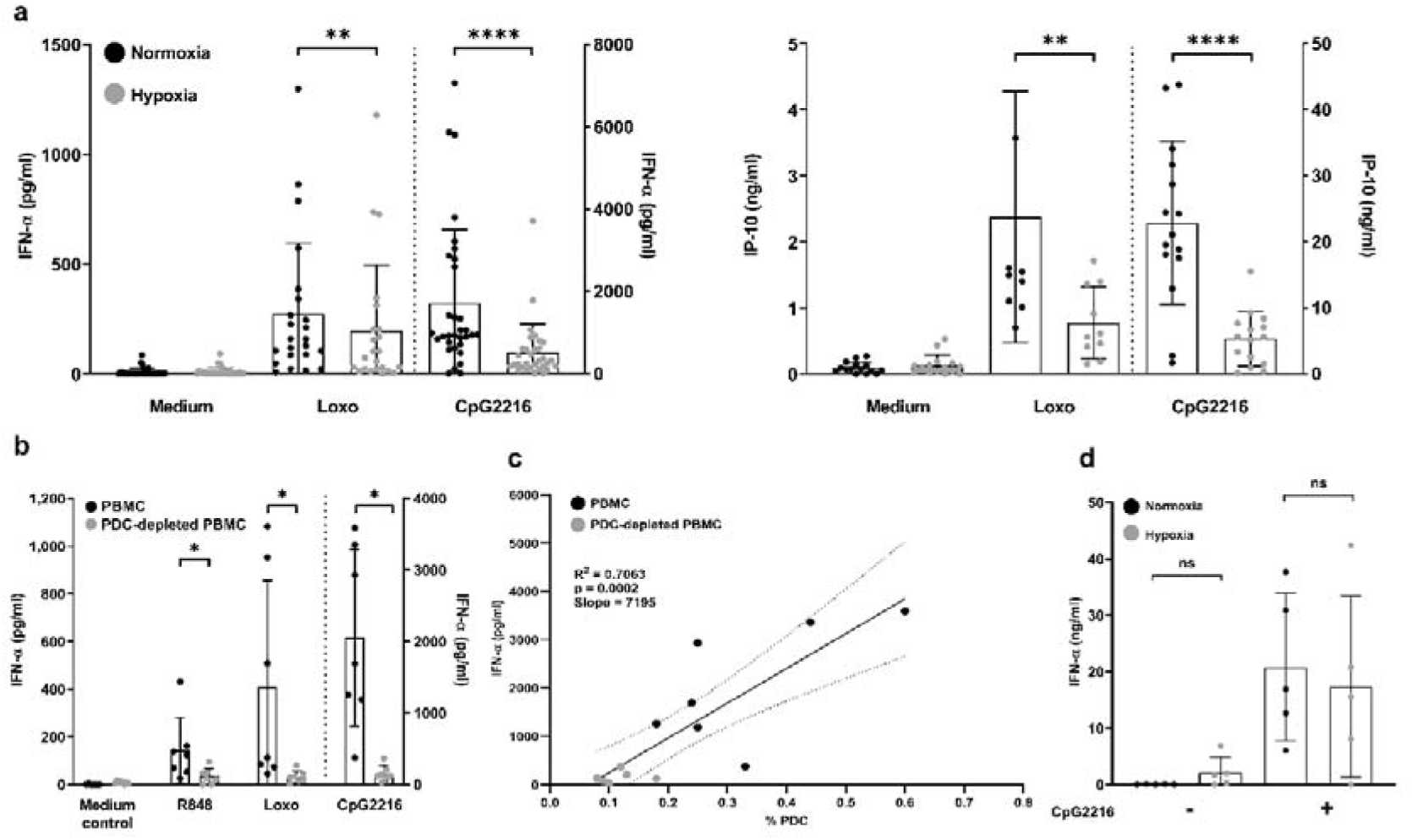
CpG2216 stimulated PDC in PBMC, but not isolated PDC secrete less IFN-α under hypoxic conditions. PBMC or PDC were adapted to normoxic or hypoxic conditions and stimulated with 0.25 µg/ml R848, 0.5 mM loxoribine or 0.5 µM CpG2216 for 20 hours, as indicated. Cytokines were measured via ELISA in supernatants. **a**. IFN-α (n = 32 donors) and IP-10 (n = 15 donors) were measured in PBMC supernatants. **b**. IFN-α was measured in supernatants of whole PBMC and PDC-depleted PBMC. Three independent experiments with n = 7 donors are shown. **c.** Linear regression of IFN-α supernatant levels after stimulation with CpG2216 (Fig. 3b) versus PDC percentage in PBMC and PDC-depleted PBMC (Fig. S3a). The line provides the slope. Dotted lines indicate 95%-confidence intervals. **d.** IFN-α was measured in supernatants of isolated PDC. Five independent experiments. n = 5 donors. Bars indicate mean ± SD. Statistics: Wilcoxon matched-pairs signed-rank tests.

Subsequent depletion of PDC from PBMC (Fig. S3a) identified PDC as the major source of IFN-α in our experimental setting and confirmed their central role in human TLR-induced IFN-α induction. Comparison of IFN-α secretion levels in the presence and absence of PDC revealed nearly complete loss of IFN-α upon stimulation with CpG2216, loxoribine or R848-stimulated PBMC (Fig. 3b). We further statistically tested whether the percentage of PDC correlated with the amount of released IFN-α ( Fig. 3c) and found the correlation to be significant (p = 0.0002, R^2^ = 0.7063). These data supported the conclusion that PDC represent the main source of hypoxia-regulated IFN-α release.

### IFN-**α** producing capacity of PDC remains intact under hypoxic conditions

Having identified PDC as the hypoxia-sensitive IFN-α producer, we next enriched PDC, adapted them to hypoxia, and stimulated with CpG2216. These experiments revealed that in some donors the ability of PDC to produce IFN-α was not affected by hypoxia (Fig. 3d). Furthermore, viability of PDC within PBMC was not affected by hypoxia as confirmed by flow cytometric analysis (Fig. S3b). This led us to the hypothesis that hypoxia-mediated changes in PDC function are indirectly supported via cellular signals originating from bystander cells in the context of PBMC, thus excluding a specific targeting of PDC by hypoxia-related factors.

### Hypoxia-mediated suppression of IFN-**α** is equally reproducible in PDC cocultures with B cells and monocytes

To better understand the cellular origin of the suppressive mechanism controlling PDC we decided to study IFN-α secretion in PDC cocultures with either CD19 positive B cells or CD14 positive monocytes. The results showed that the reduction of IFN-α in hypoxia was detectable in both settings (Fig. 4). We concluded that the signals leading to the blockage of PDC-derived IFN-α can be mediated by cells of both lymphoid and myeloid origin. Moreover, the presence of T cells was not required for the suppressive effect.

**Fig. 4:**
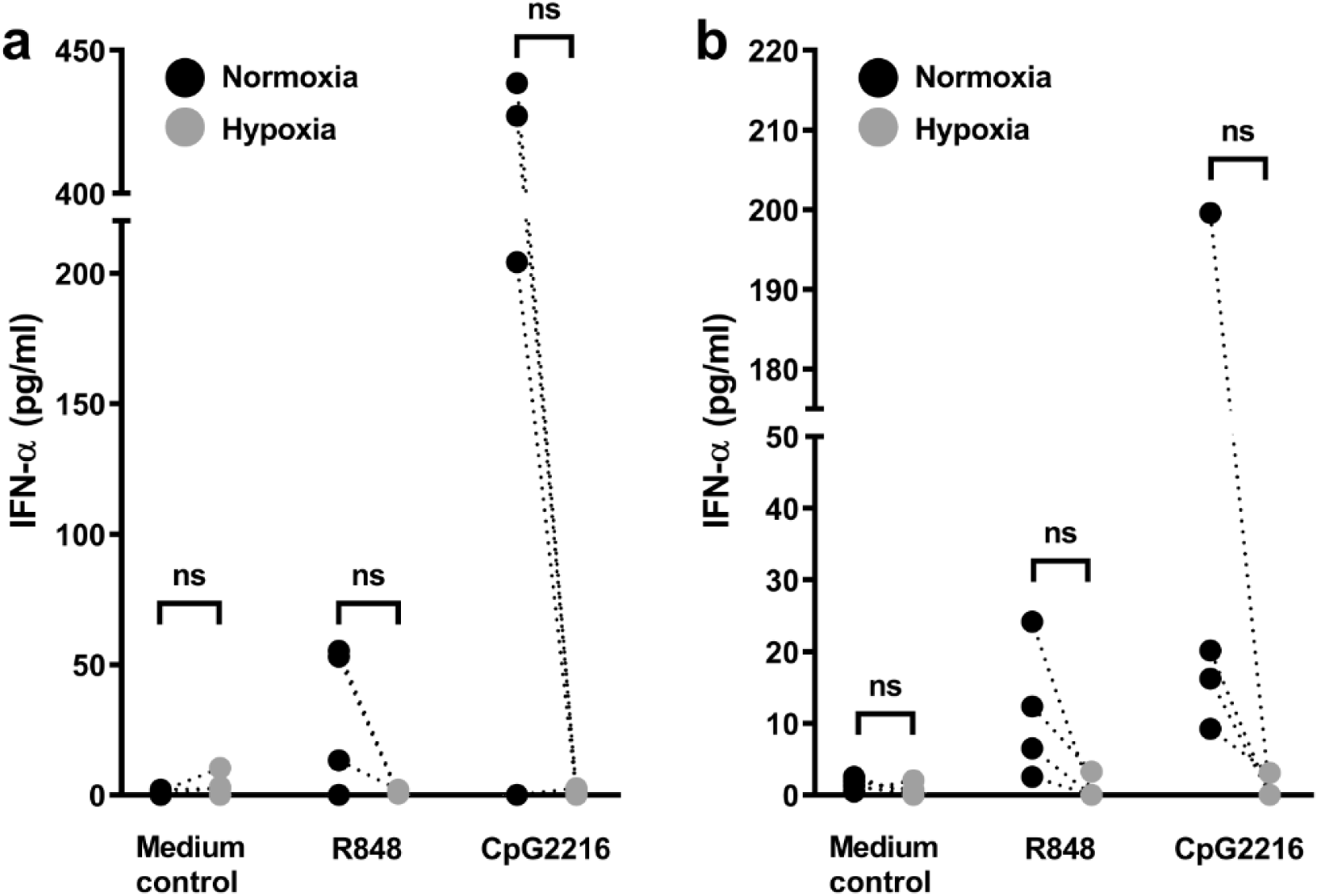
Co-incubation of PDC with B cell or monocytes is sufficient to allow for reduction of IFN-α secretion in hypoxia. PDC were co-isolated with B cells (**a**) or monocytes (**b**) from PBMC, adapted to normoxic or hypoxic conditions and stimulated with 0.5 µM CpG2216 or 0.25 µg/ml R848 for 20 hours. IFN-α was measured in supernatants. The graph shows two independent experiments with n = 4 donors. Statistical analysis was performed with Wilcoxon matched-pairs signed-rank test.

### Lactate contributes to reduction but does not block IFN-**α** secretion capacity

Next, we reasoned that soluble factors such as cytokines and metabolites secreted by hypoxic PBMC could be responsible for the suppressive effect. Since lactate, a metabolite released under hypoxic conditions from all cell types, was previously described as an immune modulator, we subsequently measured lactate in supernatants of PBMC. Cells were stimulated with CpG2216 and lactate concentrations detected ranged from 1.5 – 4 mM in hypoxia, while lactate remained below the detection limit in supernatants generated in normoxia (Fig. 5a). Lactate concentrations were further comparable in unstimulated versus CpG2216-stimulated conditions, indicating that the secretion of lactate does not require TLR stimulation.

**Fig. 5:**
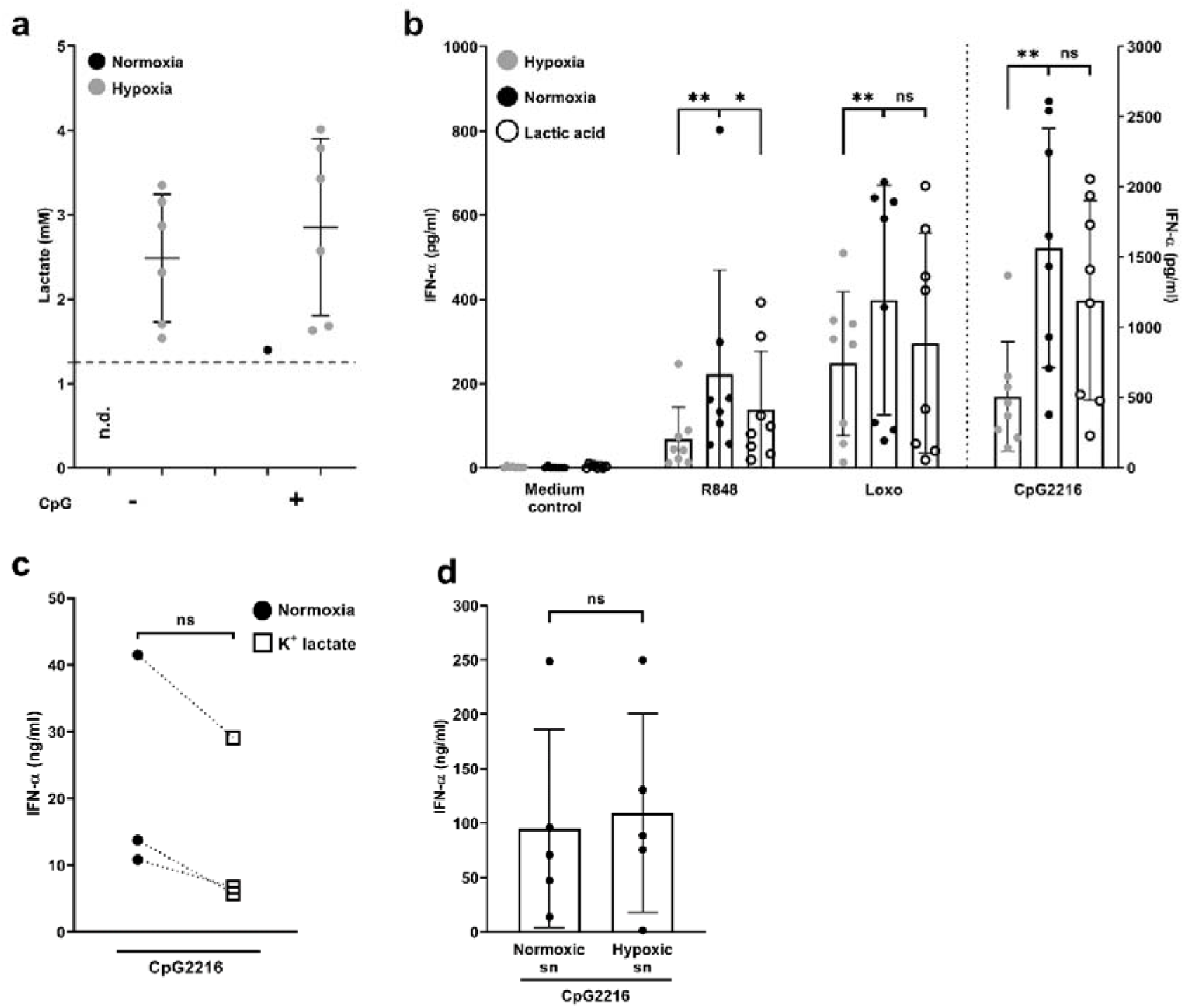
Role of lactate and purified PDC in IFN-α suppression. **a.-c.** PBMC were adapted to hypoxic conditions and stimulated with 0.5 µM CpG2216 for 20 hours. Supernatants were frozen until further use. **a.** Lactate was measured in thawed PBMC supernatants. Dotted line = detection limit (1.25 mM). Three independent experiments with n = 6 donors. **b**. PBMC were stimulated with 0.25 µg/ml R848, 0.5 mM loxoribine or 0.5 µM CpG2216 in 5 mM lactic acid. Four independent experiments with n = 8 donors. **c**. PDC were adapted to 10 mM K+ lactate and stimulated with 0.5 µM CpG2216 for or 18 ±4 hours; three independent experiments with n = 3 donors. Bars/lines indicate mean ± SD calculated with paired t-test. **d.** Isolated PDC were incubated with supernatants (sn) of (allogeneic) PBMC generated under hypoxia without stimulation for 24 hours. Subsequently, PDC were stimulated with 0.5 µM CpG2216 for 14 hours and IFN-α was measured in supernatants. n=4 independent experiments. n = 5 donors. Bars indicate mean ± SD. Statistics were calculated with paired Student’s t-test.

Since PBMC and the PDC co-cultures were cultured at higher cell density than purified PDC (2x10^6^ /ml versus 0.25x10^6^ /ml), we argued that higher amounts of lactate accumulation might be required for suppression of the IFN-α response. To assess the impact of lactate on IFN-α secretion, we treated normoxic PBMC with 5 mM lactate (dissolved lactic acid) and stimulated with CpG2216. In this setting the presence of lactate did not significantly impact IFN-α levels albeit there was a clear trend to a reduction (Fig. 5b). When we stimulated with loxoribine (TLR7) we obtained similar results: there was a trend but reduction of IFN-α was not significant (Fig. 5b). By contrast, we observed a significant reduction in IFN-α concentrations upon R848 stimulation in the presence of lactate (Fig. 5b).

To further evaluate the relevance of these findings, we incubated purified PDC with 10 mM K^+^ lactate as published by Raychaudhuri and colleagues [40]. Treatment of PDC with K^+^ lactate decreased IFN-α secretion in all donors tested. However, this reduction was only partial and not statistically significant (Fig. 5c). These data indicated that lactate can reduce IFN-I secretion but it is most likely not sufficiently active to explain the decrease of PDC-derived IFN-α under hypoxic conditions.

To further clarify the relevance of secreted factors we used supernatants generated from normoxic and hypoxic (unstimulated) PBMC to test whether accumulating lactate released from PBMC together with other mediators could suppress CpG2216- stimulated PDC-derived IFN-α secretion. The results showed that the supernatants failed to mediate IFN-α suppression (Fig. 5d). Thus, soluble factors alone might not be sufficient to mediate hypoxia-induced IFN-α blockade.

### PDC-derived IFN-**α** secretion can partially be restored by inhibition of TGF-**β** signaling

To further address the nature of the hypoxia-mediated IFN-α blockage, we analyzed whether known IFN-controlling factors are present in supernatants from PBMC in the presence and absence of hypoxia. Experiments showed that in PBMC stimulated with or without loxoribine or CpG2216 IL-10 and PGE_2_ were present in all conditions albeit slightly reduced under hypoxia (Fig. S4).

Since HIF induces TGF-β and TGF-β was previously shown to inhibit IFN-α in PDC [41], we next tried to rescue R848- and CpG2216-triggered secretion of IFN-α by pretreating PBMC with SB431542, which interferes with TGF-β signaling by inhibiting ALK5-mediated smad2 phosphorylation. The results showed that SB431542 had little effect in normoxia, but increased IFN-α secretion in hypoxia (Fig. 6, Fig. S5). Additionally, we used forskolin to assess the potential contribution of PGE_2_-mediated cAMP signaling as previously described [41]. As expected, forskolin reduced IFN-α secretion in all conditions (Fig. 6 (normalized data), Fig. S5 (IFN-α concentrations)), suggesting an additional role of constitutively secreted PGE_2_ in the control of IFN-α secretion.

**Fig. 6:**
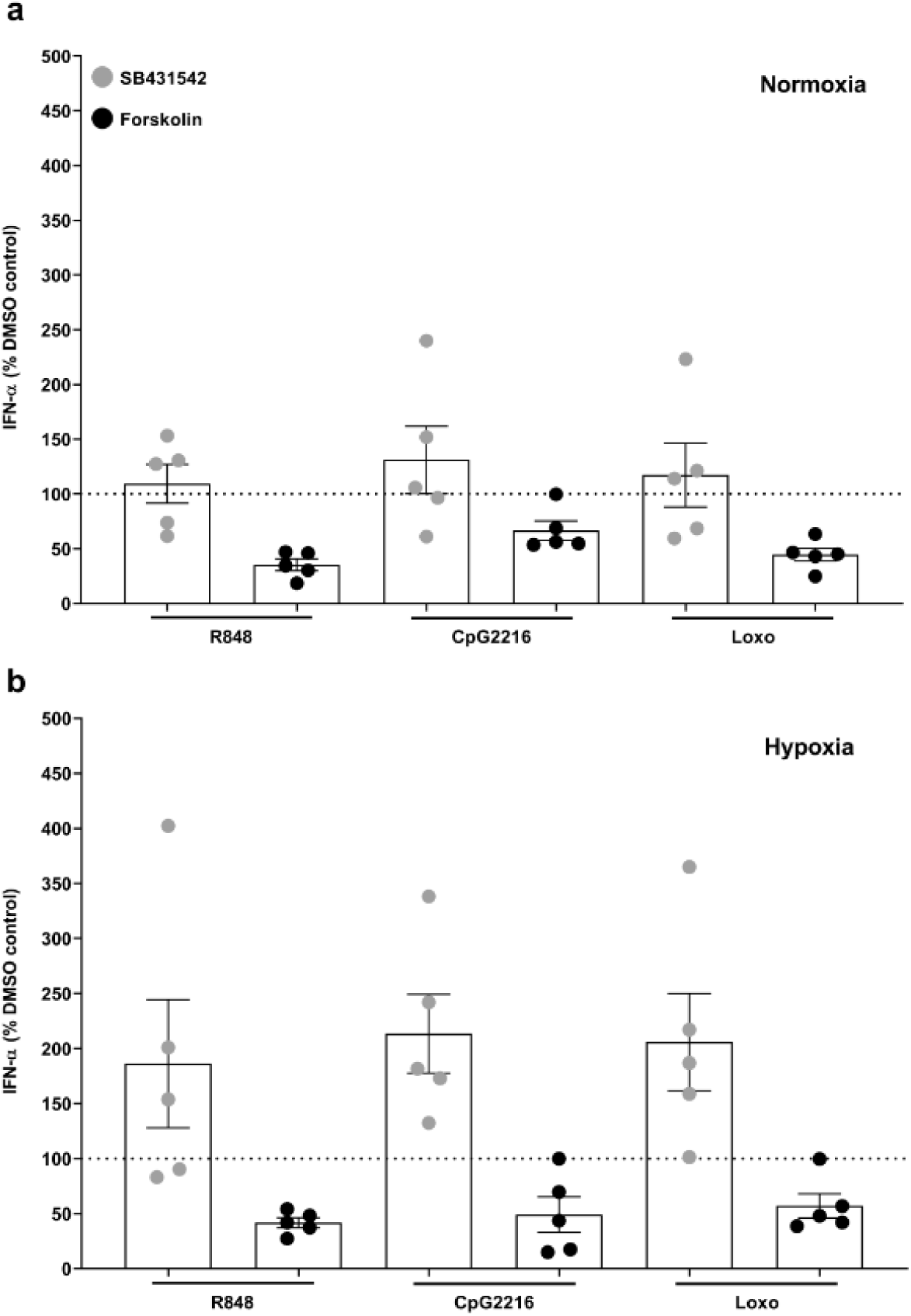
Effect of TGF-β inhibition and cAMP induction on IFN-α secretion by PBMC. PBMC were adapted to normoxia (a) or hypoxia (b) in 1 µM SB431542 (TGF-β inhibitor), 20 µM forskolin (PGE_2_ analog) or a 14.1 mM DMSO control (equivalent to condition with highest DMSO concentration: 20 µM forskolin) for 4 hours. PBMC were thereafter stimulated with 0.25 µg/ml R848, 0.5 µM CpG2216 or 0.5 mM loxoribine for 20 hours. IFN-α concentrations in supernatants were measured via ELISA. Normalized data (DMSO control = 100%, dashed line). Values obtained in three independent experiments with n = 5 donors were normalized to the respective DMSO controls. Bars indicate mean ± SEM calculated by Wilcoxon matched-pairs signed-rank test.

Taken together these data indicated that hypoxia-specific release of lactate and TGF-β could synergize with signaling induced by PGE_2_ and further mediators to promote reduction of IFN-α production.

## Discussion

Leukocytes are inevitably exposed to a broad range of oxygen concentrations. In this study, we were interested in qualitative changes in host recognition of pathogens in hypoxia. Our results demonstrate that hypoxia-mediated TLR7/9-signaling selectively regulates IFN-α responses in human PBMC (Fig. 1, Fig. 2, Fig. 3). This is an important finding because IFN-I is key to priming of innate immune cells for rapid responses to infectious pathogens [42] and a central regulator of the adaptive immune response, enabling differentiation towards Th1 profiles [43] as well as the formation of plasma cells and class switch recombination [44, 45]. This might imply that hypoxia modulates the environment to avoid strong inflammatory responses, while in the context of solid tumors modulation of IFN and PDC responses might have important implications on treatment efficacy with TLR7/9 ligands [41]. Notably, (hypoxia-induced) elevations in levels of IL-6 (Fig. 1a), a growth and differentiation factor for T and B cells [46] are characteristic of a tolerogenic environment and described in conjunction with high PGE_2_ levels and low IFN-I levels [41, 47]. By contrast preserved levels of TNF and IL-1β (Fig. 1a) indicate that hypoxia-induced modulation does not lead to unselective immune suppression.

Importantly, our data demonstrate residual IFN-α production under hypoxic conditions in PDC (observed in individual donors (Fig. 3d)), and in PBMC (Fig. 1b, c). This suggests that despite adapting to the environment PDC preserve their function. We can only speculate that previous priming or infection might desensitize PDC for hypoxia, which might be essential for reactivity of the host response. Future work will need to investigate whether hypoxia-sensitive and -resistant PDC subpopulations match previously described PDC types [48–50]. Interestingly, an earlier study in murine bone marrow-derived dendritic cells described that LPS- and CpG-mediated induction of type I interferon and of HIF-1α was more pronounced in hypoxia than in normoxia [51]. LPS-triggered type I IFN secretion was lost upon induction of myeloid cell-specific knockout of HIF-1α. Apart from the species difference that murine myeloid cells express TLR9 and respond to CpG and that the authors focused on TLR4 stimulation this study indicated that the induction of HIF-1α upon TLR stimulation is preserved under hypoxic conditions. Furthermore, LPS-induced accumulation of succinate under normoxic conditions was shown to promote stabilization of HIF-1α leading to IL-1β production in murine macrophages [52]. TLR-mediated stabilization of HIF-1α is therefore most likely not discriminative between hypoxic and normoxic conditions. However, IFN-α induction in PDC depends on IRF- 5 and NFκB [53] and the IRF5 promoter contains a hypoxia response element, which is negatively regulated by HIF-1α [54]. We assume that this mechanism contributes to loss of IFN-α production in hypoxia-sensitive PDC.

Here, we further proposed that soluble factors could serve as indirect regulators of TLR-induced IFN-α production. Notably, a study on human PDC showed that K^+^ lactate reduced the levels of IFN-α secreted by PDC in response to CpG2216 [40]. Reduced IFN-α levels were confirmed in lactate-infused mice.These authors showed that lactate exerts its effects via the cell surface receptor GPR81 as well as via cytosolic import [40]; others claimed that it acts as PHD inhibitor in normoxic conditions [55]. In our experiments, we confirmed an inhibitory effect of lactate on the IFN-I response (Fig. 5b, c) but both lactate and lactate-containing supernatants were not sufficient to fully repress IFN-α secretion (Fig. 5). We, therefore, argued that other cellular mechanisms act in synergy with lactate to enhance the suppressive effect on PDC-derived IFN-α secretion. Our data further show that chemical induction of hypoxia was not associated with lactate accumulation in the culture medium (Fig. 2c), albeit the suppressive effect on IFN-α release was reproducible with CoCl_2_ and DMOG (Fig. 2b). This suggested that stabilization of HIF-1α is an additional driver for IFN-α induction as reported in murine dendritic cells [51]. Hypoxia-induced HIF-1α expression was confirmed in whole cell lysates of PBMC (Fig. 5a).

Importantly, our data support the notion that hypoxia-modulated PDC function is highly dependent on indirect effects exerted by the cellular environment. Although supernatants of hypoxia-exposed PBMC were not suppressive, our experimental data could not exclude soluble factors as mediators of suppression because in hypoxia inhibition of TGF-β signaling unleashed IFN-α release (Fig. 6b). Forskolin-mediated suppression of IFN-α via cAMP induction suggested that PGE_2_, which was detected in supernatants of both hypoxic and normoxic PBMC (Fig. S4b), most likely contributes to the suppressory effect independent of O_2_ saturation (Fig. 6 and Fig. S5). Thus, suppression of IFN-α arises from combined suppression by multiple factors present under hypoxic conditions.

At this point we cannot exclude that cell contact or vicinity might be needed to facilitate the inhibitory effects of cellular metabolites produced in minute amounts and/or unstable in the supernatants. Downregulation of PDC-derived IFN-α production is, thus, subject to fine adjustment and a temporary event. Its main purpose could be to minimize the amplificatory potential of IFN-I, while total blockade of IFN-I release would endanger the PDĆs role in surveillance of invading pathogens.

Taken together, this study demonstrates that hypoxia-mediated regulation of the innate immune cell function occurs in a highly selective manner. Selective blockade of IFN-I-mediated amplification loops is an important mechanism to avoid excessive immune activation and tissue damage. In this respect, our study highlights the central role of PDC as hypoxia-sensitive target cell. However, future work is needed to unravel the exact cellular and molecular mechanisms underlying this highly sophisticated immune control. Moreover, the comparison of stimulatory conditions that activate PDC alone or together with monocytes indicates that suppression depends on the activation status of the cellular environment and concentrations of IFN-regulatory factors. Thus, inhibition could be reverted by stimulation of the bystander cells, which would in turn be key to reinstall a proinflammatory immune response when needed.

## Supporting information

Supplemental figures Lenkewitz et al.

## Acknowledgments

We would like to thank all lab colleagues that supported our work.

## Funding

This project was financed by intramural funding of the Paul-Ehrlich-Institut.

## Author contributions

AL and IBD designed the study. AL, FH, NI, OT and IBB performed experiments. AL, IBD, FH, NI, OT and IBB analyzed the data. AL and IBD designed the figures, interpreted the data and wrote the manuscript.

## Notes

### Competing Interest Statement

The authors have declared no competing interest.

## References

1 Falchuk, K. H., Goetzl, E. J. and Kulka, J. P., Respiratory gases of synovial fluids. An approach to synovial tissue circulatory-metabolic imbalance in rheumatoid arthritis. The American journal of medicine. 1970. 49: 223–231.

2 Abbott, R. K., Thayer, M., Labuda, J., Silva, M., Philbrook, P., Cain, D. W. and Kojima, H. et al., Germinal Center Hypoxia Potentiates Immunoglobulin Class Switch Recombination. Journal of immunology (Baltimore, Md. 1950). 2016. 197: 4014–4020.

3 Jantsch, J. and Schödel, J., Hypoxia and hypoxia-inducible factors in myeloid cell-driven host defense and tissue homeostasis. Immunobiology. 2015. 220: 305–314.

4 Semenza, G. L., Nejfelt, M. K., Chi, S. M. and Antonarakis, S. E., Hypoxia-inducible nuclear factors bind to an enhancer element located 3’ to the human erythropoietin gene. Proceedings of the National Academy of Sciences of the United States of America. 1991. 88: 5680–5684.

5 Wang, G. L., Jiang, B. H., Rue, E. A. and Semenza, G. L., Hypoxia-inducible factor 1 is a basic-helix-loop-helix-PAS heterodimer regulated by cellular O2 tension. Proceedings of the National Academy of Sciences of the United States of America. 1995. 92: 5510–5514.

6 Maxwell, P. H., Wiesener, M. S., Chang, G. W., Clifford, S. C., Vaux, E. C., Cockman, M. E. and Wykoff, C. C. et al., The tumour suppressor protein VHL targets hypoxia-inducible factors for oxygen-dependent proteolysis. Nature. 1999. 399: 271–275.

7 Ivan, M., Kondo, K., Yang, H., Kim, W., Valiando, J., Ohh, M. and Salic, A. et al., HIFalpha targeted for VHL-mediated destruction by proline hydroxylation: implications for O2 sensing. Science (New York, N.Y.). 2001. 292: 464–468.

8 Jaakkola, P., Mole, D. R., Tian, Y. M., Wilson, M. I., Gielbert, J., Gaskell, S. J. and Kriegsheim, A. von, et al., Targeting of HIF-alpha to the von Hippel-Lindau ubiquitylation complex by O2- regulated prolyl hydroxylation. Science (New York, N.Y.). 2001. 292: 468–472.

9 Lee, P., Chandel, N. S. and Simon, M. C., Cellular adaptation to hypoxia through hypoxia inducible factors and beyond. Nature reviews. Molecular cell biology. 2020. 21: 268–283.

10 Wenger, R. H., Kurtcuoglu, V., Scholz, C. C., Marti, H. H. and Hoogewijs, D., Frequently asked questions in hypoxia research. Hypoxia (Auckland, N.Z.). 2015. 3: 35–43.

11 Keeley, T. P. and Mann, G. E., Defining Physiological Normoxia for Improved Translation of Cell Physiology to Animal Models and Humans. Physiological reviews. 2019. 99: 161–234.

12 Carreau, A., El Hafny-Rahbi, B., Matejuk, A., Grillon, C. and Kieda, C., Why is the partial oxygen pressure of human tissues a crucial parameter? Small molecules and hypoxia. Journal of cellular and molecular medicine. 2011. 15: 1239–1253.

13 Krzywinska, E. and Stockmann, C., Hypoxia, Metabolism and Immune Cell Function. Biomedicines. 2018. 6.

14 Eltzschig, H. K. and Carmeliet, P., Hypoxia and inflammation. The New England journal of medicine. 2011. 364: 656–665.

15 Nizet, V. and Johnson, R. S., Interdependence of hypoxic and innate immune responses. Nature reviews. Immunology. 2009. 9: 609–617.

16 Pham, K., Parikh, K. and Heinrich, E. C., Hypoxia and Inflammation: Insights From High-Altitude Physiology. Frontiers in physiology. 2021. 12: 676782.

17 Colgan, S. P., Furuta, G. T. and Taylor, C. T., Hypoxia and Innate Immunity: Keeping Up with the HIFsters. Annual review of immunology. 2020. 38: 341–363.

18 Stothers, C. L., Luan, L., Fensterheim, B. A. and Bohannon, J. K., Hypoxia-inducible factor-1α regulation of myeloid cells. Journal of molecular medicine. 2018. 96: 1293–1306.

19 Frede, S., Stockmann, C., Freitag, P. and Fandrey, J., Bacterial lipopolysaccharide induces HIF-1 activation in human monocytes via p44/42 MAPK and NF-kappaB. The Biochemical journal. 2006. 396: 517–527.

20 Spirig, R., Djafarzadeh, S., Regueira, T., Shaw, S. G., Garnier, C. von, Takala, J. and Jakob, S. M., et al., Effects of TLR agonists on the hypoxia-regulated transcription factor HIF-1alpha and dendritic cell maturation under normoxic conditions. PLoS ONE. 2010. 5: e0010983.

21 Schaffer, K. and Taylor, C. T., The impact of hypoxia on bacterial infection. The FEBS journal. 2015. 282: 2260–2266.

22 Fitzpatrick, S. F., Tambuwala, M. M., Bruning, U., Schaible, B., Scholz, C. C., Byrne, A. and O’Connor, A. et al., An intact canonical NF-κB pathway is required for inflammatory gene expression in response to hypoxia. Journal of immunology (Baltimore, Md. 1950). 2011. 186: 1091–1096.

23 Naldini, A. and Carraro, F., Hypoxia modulates cyclin and cytokine expression and inhibits peripheral mononuclear cell proliferation. Journal of cellular physiology. 1999. 181: 448–454.

24 Naldini, A., Carraro, F., Silvestri, S. and Bocci, V., Hypoxia affects cytokine production and proliferative responses by human peripheral mononuclear cells. Journal of cellular physiology. 1997. 173: 335–342.

25 Cho, S. H., Raybuck, A. L., Blagih, J., Kemboi, E., Haase, V. H., Jones, R. G. and Boothby, M. R., Hypoxia-inducible factors in CD4+ T cells promote metabolism, switch cytokine secretion, and T cell help in humoral immunity. Proceedings of the National Academy of Sciences of the United States of America. 2019. 116: 8975–8984.

26 Yang, M., Ma, C., Liu, S., Sun, J., Shao, Q., Gao, W. and Zhang, Y. et al., Hypoxia skews dendritic cells to a T helper type 2-stimulating phenotype and promotes tumour cell migration by dendritic cell-derived osteopontin. Immunology. 2009. 128: e237–49.

27 Wang, Q., Liu, C., Zhu, F., Liu, F., Zhang, P., Guo, C. and Wang, X. et al., Reoxygenation of hypoxia-differentiated dentritic cells induces Th1 and Th17 cell differentiation. Molecular immunology. 2010. 47: 922–931.

28 Roman, J., Rangasamy, T., Guo, J., Sugunan, S., Meednu, N., Packirisamy, G. and Shimoda, L. A. et al., T-cell activation under hypoxic conditions enhances IFN-gamma secretion. American journal of respiratory cell and molecular biology. 2010. 42: 123–128.

29 Kim, S. Y., Choi, Y. J., Joung, S. M., Lee, B. H., Jung, Y.-S. and Lee, J. Y., Hypoxic stress up- regulates the expression of Toll-like receptor 4 in macrophages via hypoxia-inducible factor. Immunology. 2010. 129: 516–524.

30 Lahat, N., Rahat, M. A., Ballan, M., Weiss-Cerem, L., Engelmayer, M. and Bitterman, H., Hypoxia reduces CD80 expression on monocytes but enhances their LPS-stimulated TNF-alpha secretion. Journal of leukocyte biology. 2003. 74: 197–205.

31 Hoque, R., Farooq, A., Ghani, A., Gorelick, F. and Mehal, W. Z., Lactate reduces liver and pancreatic injury in Toll-like receptor- and inflammasome-mediated inflammation via GPR81- mediated suppression of innate immunity. Gastroenterology. 2014. 146: 1763–1774.

32 Errea, A., Cayet, D., Marchetti, P., Tang, C., Kluza, J., Offermanns, S., Sirard, J.-C. and Rumbo, M., Lactate inhibits the pro-inflammatory response and metabolic reprogramming in murine macrophages in a GPR81-independent manner. Universitätsbibliothek Johann Christian Senckenberg, Frankfurt am Main 2016.

33 Ratter, J. M., Rooijackers, H. M. M., Hooiveld, G. J., Hijmans, A. G. M., Galan, B. E. de, Tack, C. J. and Stienstra, R., In vitro and in vivo Effects of Lactate on Metabolism and Cytokine Production of Human Primary PBMCs and Monocytes. Frontiers in immunology. 2018. 9: 2564.

34 Dietl, K., Renner, K., Dettmer, K., Timischl, B., Eberhart, K., Dorn, C. and Hellerbrand, C. et al., Lactic Acid and Acidification Inhibit TNF Secretion and Glycolysis of Human Monocytes. The Journal of Immunology. 2010. 184: 1200–1209.

35 Gottfried, E., Kunz-Schughart, L. A., Ebner, S., Mueller-Klieser, W., Hoves, S., Andreesen, R., Mackensen, A. and Kreutz, M., Tumor-derived lactic acid modulates dendritic cell activation and antigen expression. Blood. 2006. 107: 2013–2021.

36 Bekeredjian-Ding, I., Roth, S. I., Gilles, S., Giese, T., Ablasser, A., Hornung, V., Endres, S. and Hartmann, G., T cell-independent, TLR-induced IL-12p70 production in primary human monocytes. Journal of immunology (Baltimore, Md. 1950). 2006. 176: 7438–7446.

37 Hornung, V., Rothenfusser, S., Britsch, S., Krug, A., Jahrsdörfer, B., Giese, T., Endres, S. and Hartmann, G., Quantitative expression of toll-like receptor 1-10 mRNA in cellular subsets of human peripheral blood mononuclear cells and sensitivity to CpG oligodeoxynucleotides. Journal of immunology (Baltimore, Md. 1950). 2002. 168: 4531–4537.

38 Kerkmann, M., Rothenfusser, S., Hornung, V., Towarowski, A., Wagner, M., Sarris, A. and Giese, T. et al., Activation with CpG-A and CpG-B oligonucleotides reveals two distinct regulatory pathways of type I IFN synthesis in human plasmacytoid dendritic cells. Journal of immunology (Baltimore, Md. 1950). 2003. 170: 4465–4474.

39 Bekeredjian-Ding, I. B., Berkeredjian-Ding, I. B., Wagner, M., Hornung, V., Giese, T., Schnurr, M., Endres, S. and Hartmann, G., Plasmacytoid dendritic cells control TLR7 sensitivity of naive B cells via type I IFN. Journal of immunology (Baltimore, Md. 1950). 2005. 174: 4043–4050.

40 Raychaudhuri, D., Bhattacharya, R., Sinha, B. P., Liu, C. S. C., Ghosh, A. R., Rahaman, O. and Bandopadhyay, P. et al., Lactate Induces Pro-tumor Reprogramming in Intratumoral Plasmacytoid Dendritic Cells. Frontiers in immunology. 2019. 10: 1878.

41 Bekeredjian-Ding, I., Schäfer, M., Hartmann, E., Pries, R., Parcina, M., Schneider, P. and Giese, T. et al., Tumour-derived prostaglandin E2 and transforming growth factor-&bgr; synergize to inhibit plasmacytoid dendritic cell-derived interferon-&agr;. Immunology an official journal of the British Society for Immunology the journal of cells, molecules, systems and technologies. 2009. 128: 439–450.

42 McNab, F., Mayer-Barber, K., Sher, A., Wack, A. and O’Garra, A., Type I interferons in infectious disease. Nature reviews. Immunology. 2015. 15: 87–103.

43 Brinkmann, V., Geiger, T., Alkan, S. and Heusser, C. H., Interferon alpha increases the frequency of interferon gamma-producing human CD4+ T cells. The Journal of experimental medicine. 1993. 178: 1655–1663.

44 Jego, G., Palucka, A. K., Blanck, J.-P., Chalouni, C., Pascual, V. and Banchereau, J., Plasmacytoid dendritic cells induce plasma cell differentiation through type I interferon and interleukin 6. Immunity. 2003. 19: 225–234.

45 Douagi, I., Gujer, C., Sundling, C., Adams, W. C., Smed-Sörensen, A., Seder, R. A., Karlsson Hedestam, G. B. and Loré, K., Human B cell responses to TLR ligands are differentially modulated by myeloid and plasmacytoid dendritic cells. Journal of immunology (Baltimore, Md. 1950). 2009. 182: 1991–2001.

46 Murphy, K. P., Janeway, C., Travers, P., Walport, M., Mowat, A. and Weaver, C. T., Janeway’s immunobiology, 8th edn. Garland Science, London 2012.

47 Bailly, S., Ferrua, B., Fay, M. and Gougerot-Pocidalo, M. A., Differential regulation of IL 6, IL 1 A, IL 1 beta and TNF alpha production in LPS-stimulated human monocytes: role of cyclic AMP. Cytokine. 1990. 2: 205–210.

48 Alculumbre, S. G., Diversification of human plasmacytoid predendritic cells in response to a single stimulus. Nature immunology. 2017: 1–13.

49 Thiel, A., Kesselring, R., Pries, R., Wittkopf, N., Puzik, A. and Wollenberg, B., Plasmacytoid dendritic cell subpopulations in head and neck squamous cell carcinoma. Oncology reports. 2011. 26: 615–620.

50 Bauer, J., Dress, R. J., Schulze, A., Dresing, P., Ali, S., Deenen, R., Alferink, J. and Scheu, S., Cutting Edge: IFN-β Expression in the Spleen Is Restricted to a Subpopulation of Plasmacytoid Dendritic Cells Exhibiting a Specific Immune Modulatory Transcriptome Signature. Journal of immunology (Baltimore, Md. 1950). 2016. 196: 4447–4451.

51 Wobben, R., Hüsecken, Y., Lodewick, C., Gibbert, K., Fandrey, J. and Winning, S., Role of hypoxia inducible factor-1α for interferon synthesis in mouse dendritic cells. Biological chemistry. 2013. 394: 495–505.

52 Tannahill, G. M., Curtis, A. M., Adamik, J., Palsson-McDermott, E. M., McGettrick, A. F., Goel, G. and Frezza, C. et al., Succinate is an inflammatory signal that induces IL-1β through HIF-1α. Nature. 2013. 496: 238–242.

53 Steinhagen, F., McFarland, A. P., Rodriguez, L. G., Tewary, P., Jarret, A., Savan, R. and Klinman, D. M., IRF-5 and NF-κB p50 co-regulate IFN-β and IL-6 expression in TLR9-stimulated human plasmacytoid dendritic cells. European journal of immunology. 2013. 43: 1896–1906.

54 Peng, T., Du, S.-Y., Son, M. and Diamond, B., HIF-1α is a negative regulator of interferon regulatory factors: Implications for interferon production by hypoxic monocytes. Proceedings of the National Academy of Sciences of the United States of America. 2021. 118.

55 Saedeleer, C. J. de, Copetti, T., Porporato, P. E., Verrax, J., Feron, O. and Sonveaux, P., Lactate activates HIF-1 in oxidative but not in Warburg-phenotype human tumor cells. PLoS ONE. 2012. 7: e46571.

