## Supplemental figures Lenkewitz et al. for "Hypoxia-mediated fine-tuning of the TLR7/9-triggered human PDC-derived IFN-α response is mediated by combined cellular and soluble IFN-regulators"

**Supplementary Figures**

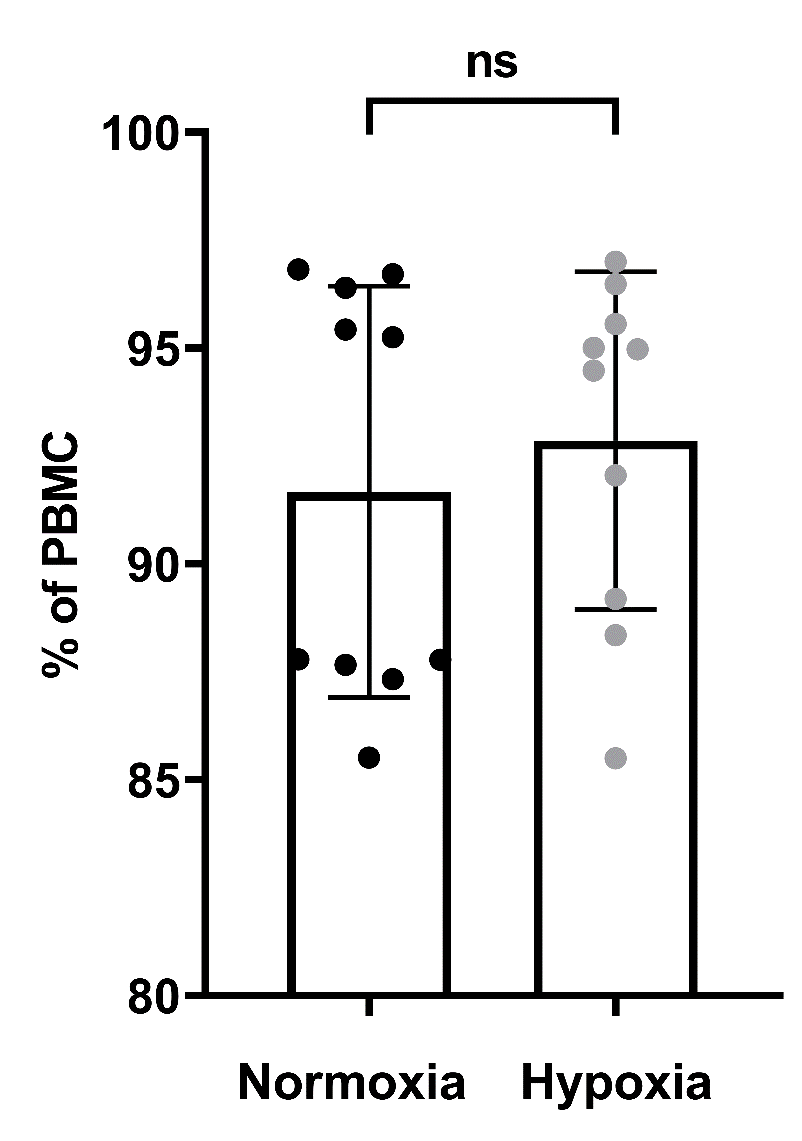

**Fig. S1: Viability of PBMC.** PBMC were adapted to hypoxic conditions and incubated for 20 hours without stimulation. Viability of PBMC was measured via flow cytometry obtained in n = 10 donors. Bars indicate mean ± SD calculated with Wilcoxon matched-pairs signed-rank test.

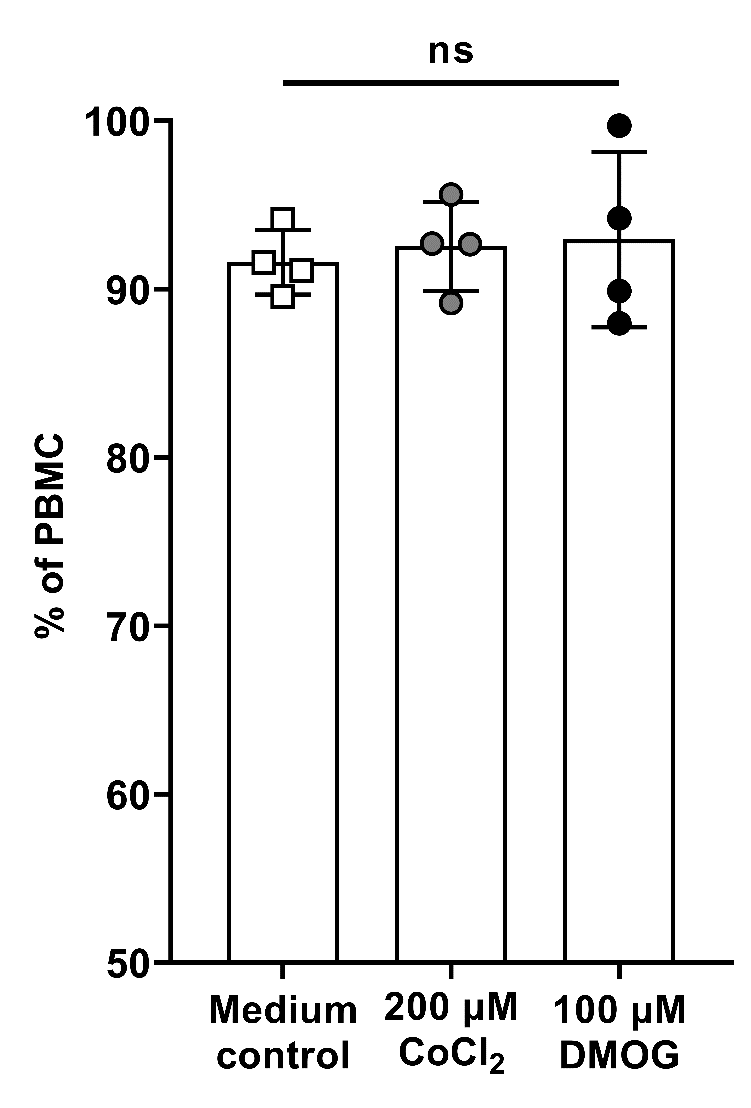

**Fig. S2: Viability of PBMC and lactate levels in PBMC supernatants.** The percentage of viable PBMC (pool of unstimulated PBMC and PBMC stimulated with 0.25 µg/ml R848, 0.5 µM CpG2216 and 0.5 mM loxoribine) was measured after 20 hours of incubation in chemical hypoxia inducers via flow cytometry. Two independent experiments from n = 4 donors. Bars indicate mean + SD calculated with paired t-test (each condition versus medium control).

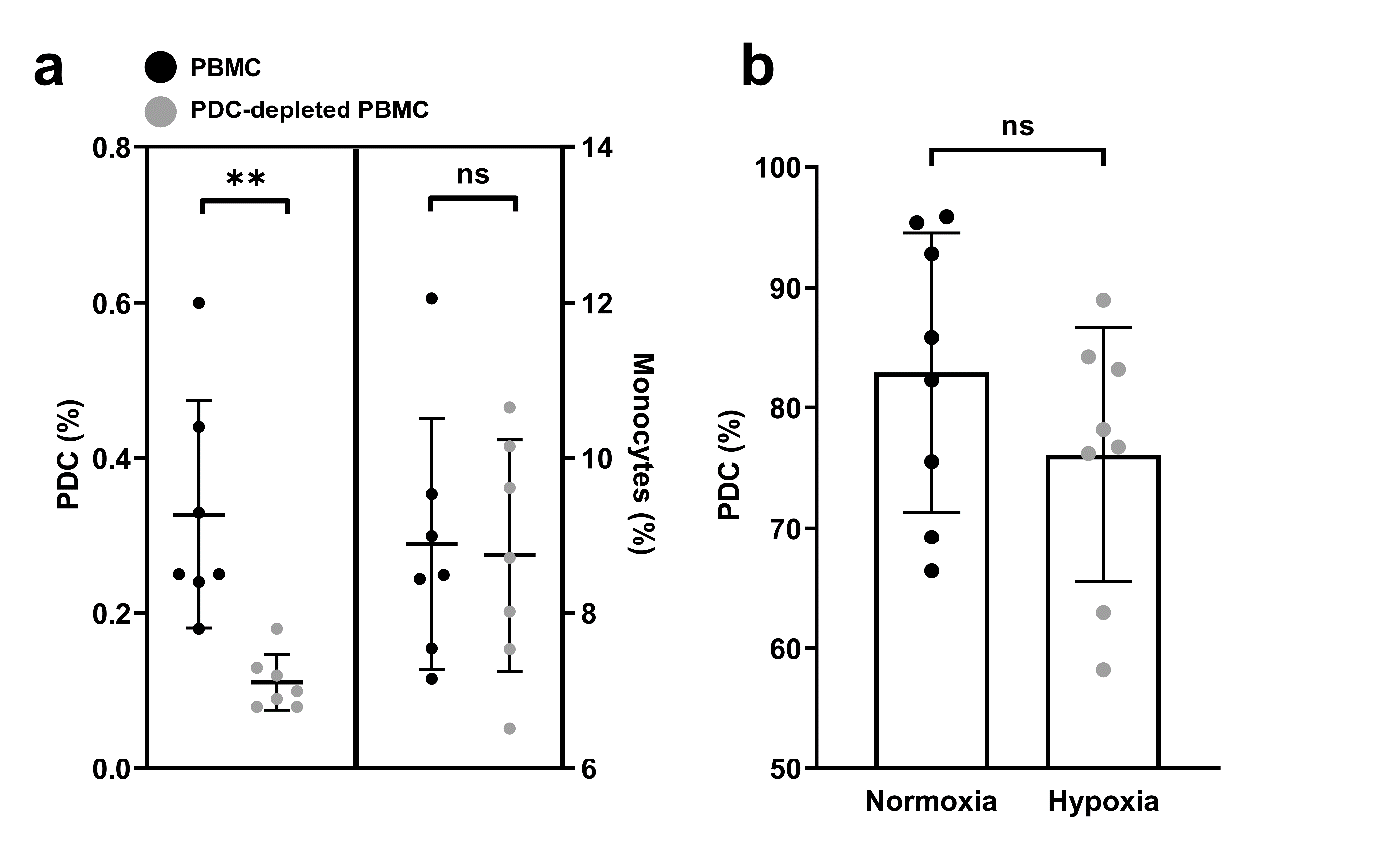

**Fig. S3: PDC are the primary IFN-α source in PBMC.** **a.** PDC depletion was controlled by flow cytometric analysis of remaining PDC. PDC and monocytes were quantified in PDC-depleted PBMC. Three independent experiments with n = 7 donors are shown. Bars indicate mean ± SD calculated with paired t-test. **b.** Percentage of viable PDC within PBMC was measured after 20 hours of stimulation with CpG2216 via flow cytometry in n = 8 donors from four independent experiments. Bars indicate mean ± SD analyzed with paired Student´s t-test.

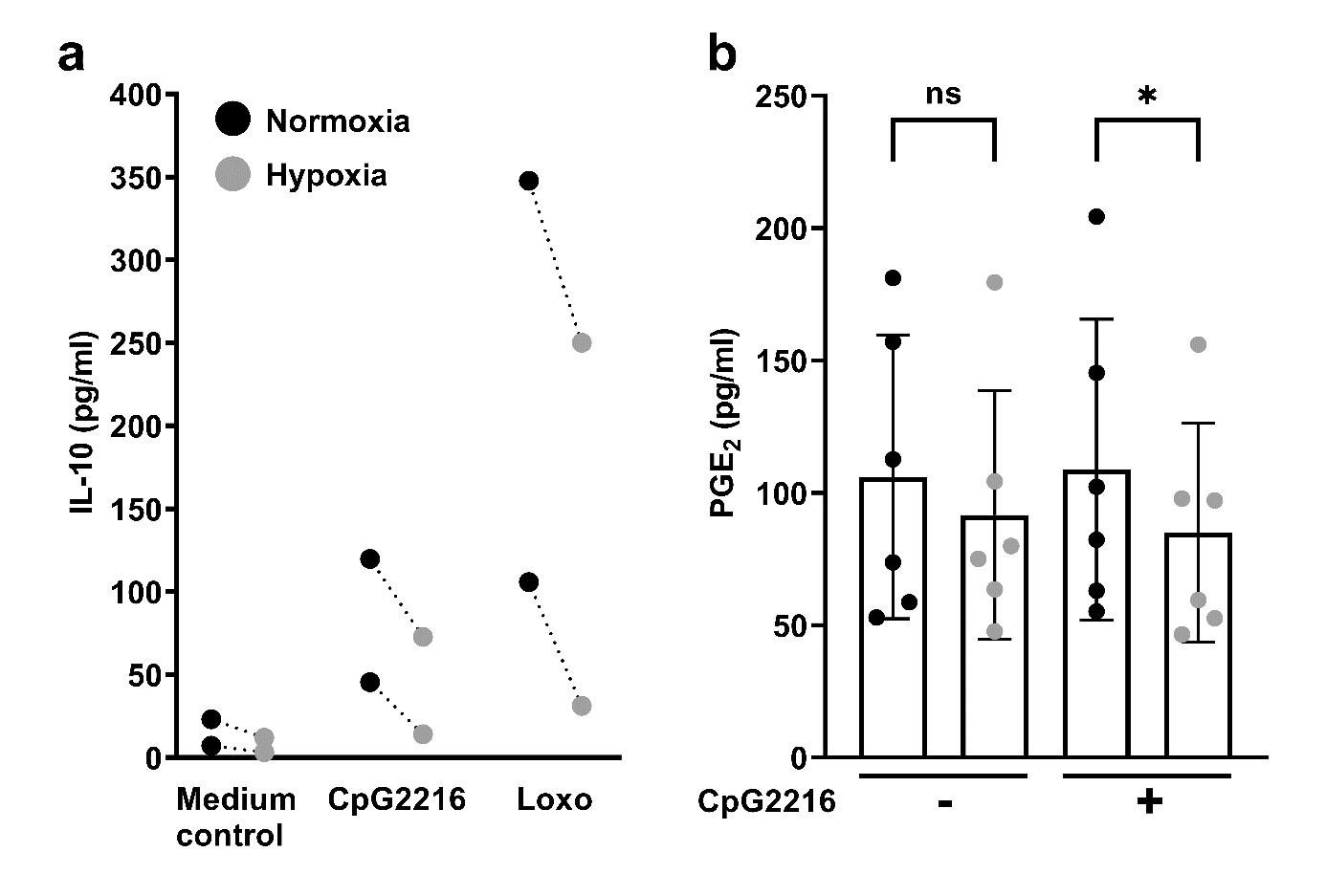

**Fig. S4: IL-10 and PGE_2_ levels in hypoxic PBMC.** PBMC were adapted to hypoxic conditions and stimulated with 0.5 µM CpG2216 or 0.5 mM loxoribine for 20 hours. **a.** IL-10 was measured via Multiplex assay. One experiment; n = 2 donors. **b.** PGE_2_ was measured via ELISA. Three independent experiments with n = 6 donors are shown. Bars indicate mean ± SD calculated with paired Student´s t-test.

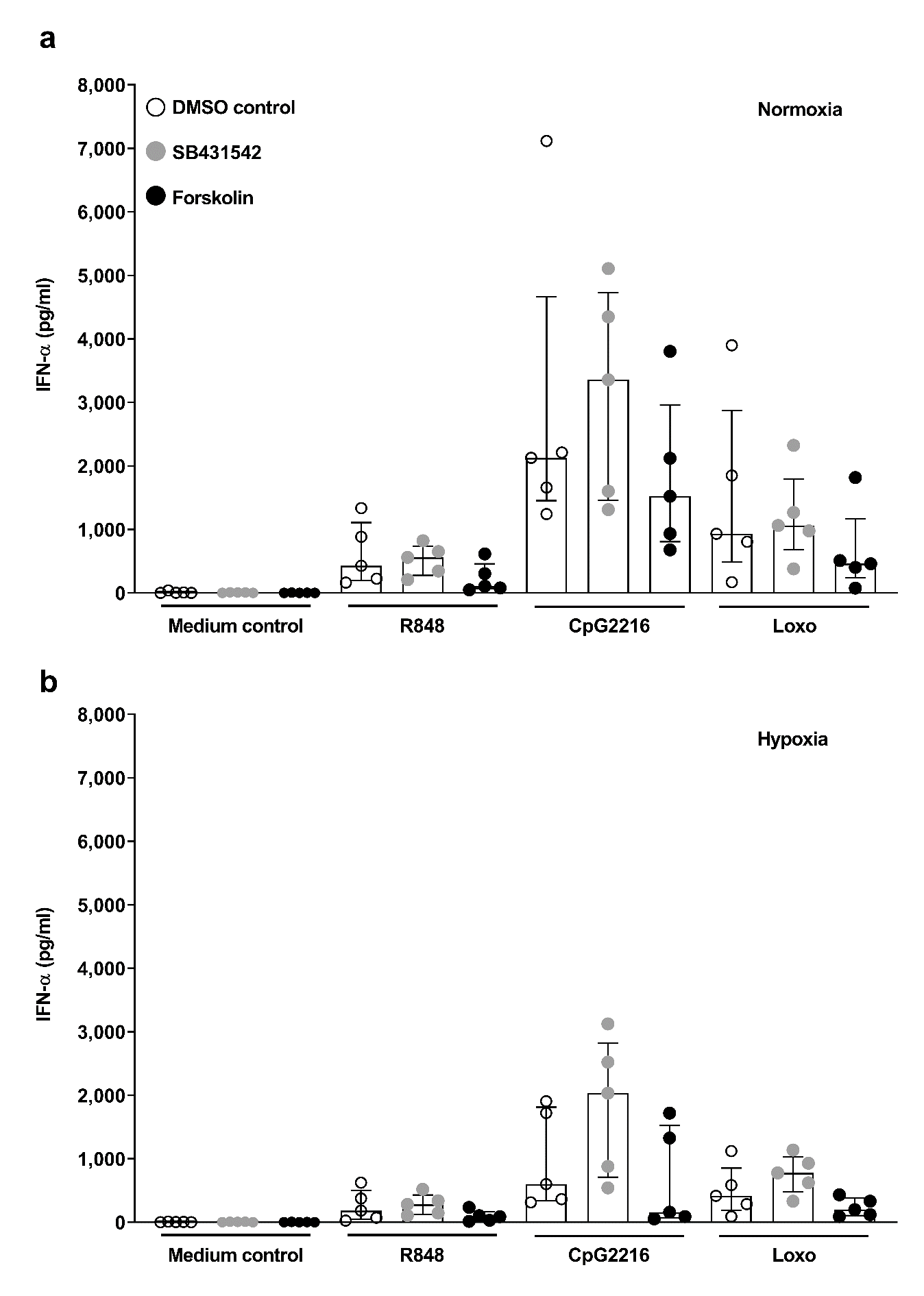

**Fig. S5:** **Effect of TGF-β inhibitor and cAMP inducer forskolin on IFN-α secretion by PBMC.** PBMC were adapted to normoxia (a) or hypoxia (b) in 1 µM SB431542, 20 µM forskolin or a 14.1 mM DMSO control for 4 hours. PBMC were then stimulated with 0.25 µg/ml R848, 0.5 µM CpG2216 or 0.5 mM loxoribine for 20 hours. IFN-α concentrations in supernatants were measured via ELISA. are IFN-α concentrations of three independent experiments with n = 5 donors are given in ng/ml. Bars indicate median ± interquartile range assessed with Wilcoxon matched-pairs signed-rank test.
